# The voltage sensing phosphatase (VSP) localizes to the apical membrane of kidney tubule epithelial cells

**DOI:** 10.1101/483743

**Authors:** Wil Ratzan, Vamseedhar Rayaprolu, Scott E. Killian, Roger Bradley, Susy C. Kohout

**Affiliations:** Department of Cell Biology and Neuroscience, Montana State University, Bozeman, MT, USA

## Abstract

Voltage-sensing phosphatases (VSPs) are transmembrane proteins that couple changes in membrane potential to hydrolysis of inositol signaling lipids. VSPs catalyze the dephosphorylation of phosphatidylinositol phosphates (PIPs) that regulate diverse aspects of cell membrane physiology including cell division, growth and migration. VSPs are highly conserved among chordates, and their RNA transcripts have been detected in the adult and embryonic stages of frogs, fish, chickens, mice and humans. However, the subcellular localization and biological function of VSP remains unknown. Using reverse transcriptase-PCR (RT-PCR), we show that both *Xenopus laevis* VSP (Xl-VSP1 and Xl-VSP2) mRNAs are expressed in early embryos, suggesting that both Xl-VSPs are involved in early tadpole development. To understand which embryonic tissues express Xl-VSP mRNA, we used *in situ* hybridization (ISH) and found Xl-VSP mRNA in both the brain and kidney of NF stage 32-36 embryos. By Western blot analysis with a VSP antibody, we show increasing levels of Xl-VSP protein in the developing embryo, and by immunohistochemistry (IHC), we demonstrate that Xl-VSP protein is specifically localized to the apical membrane of both embryonic and adult kidney tubules. We further characterized the catalytic activity of both Xl-VSP homologs and found that while Xl-VSP1 catalyzes 3- and 5-phosphate removal, Xl-VSP2 is a less efficient 3-phosphatase with different substrate specificity. Our results suggest that Xl-VSP1 and Xl-VSP2 serve different functional roles and that VSPs are an integral component of voltage-dependent PIP signaling pathways during vertebrate kidney tubule development and function.

## Introduction

Phosphatidylinositol phosphates (PIPs) are lipid second messengers involved in almost all facets of cell biology, including differentiation, proliferation, migration, and polarity (1,2). Many human diseases are linked to mutations in PIP-modifying enzymes, including cancer, peripheral neuropathy, stroke, bipolar disorder, autism and developmental disorders (3-6). As a result, PIP kinases and phosphatases have been extensively studied to understand their roles in these diverse cellular processes and are increasingly viewed as potential therapeutic targets (7-9). Here, we focus on an exceptional member of the PIP-modifying family of enzymes, the voltage-sensing phosphatase (VSP).

VSP is a unique PIP-modifying enzyme whose activity is regulated through a voltage sensing domain (VSD) (10). VSDs are composed of four transmembrane helices with the fourth helix (called S4) containing arginines responsible for sensing changes in the electrical membrane potential. In response to membrane depolarizations, the S4 helix changes conformation leading to activation of the cytosolic phosphatase domain (PD), which is homologous to PTEN (phosphatase and tensin homolog deleted on chromosome 10) (10), a well characterized lipid phosphatase. Once VSP is activated, its PD dephosphorylates both the 5- and 3-phosphates from PIPs (10-14). As a result, VSPs provide a direct connection between the electrical signaling and PIP signaling pathways. While electrical signaling pathways are most often discussed in terms of neurons, all cells maintain an ionic gradient that creates a membrane potential. This electrochemical force is utilized to initiate essential cell-signaling functions. For example, pancreatic beta cells use their membrane potential to respond to increasing glucose concentrations by activating ATP-sensitive potassium channels and voltage-gated calcium channels to release insulin (15). Renal tubules also use membrane potentials, specifically activating different potassium channels for regulating cell volume, potassium secretion and tubuloglomerular feedback (16). Since many ion channels are regulated by PIPs (17-19), VSPs are likely to play a role in modulating ion channel permeability and their subsequent electrical signaling when channels and VSP are co-expressed.

In line with the idea that VSP serves a fundamental physiological role, VSP orthologs have been identified in several tissues across diverse phyla, including ascidians, newts, salamanders, zebrafish, chicken, amphibians, mice, and humans (10,11,20–26). In particular, VSP mRNA transcripts have been found in the adult nervous systems of sea squirts and the brains of frogs, mice and humans (20,22,27,28) as well as in kidney, stomach, heart, testis and ovary (20,22–25,27–30). In addition to adult tissues, VSP transcripts have been detected in embryonic tissues such as the kidney and eye of zebrafish (31), the kidney, brain and stomach of chicks (21,30), and the brain, spinal cord and eye of mice (25,27,32). The majority of studies investigating VSP localization focused on measuring mRNA expression, with a few studies reporting cellular localization of VSP protein. The first report of the sea squirt VSP (*Ciona intestinalis* VSP, Ci-VSP) found protein expression in sperm tails (10). In chicks, *Gallus gallus* VSP (Gg-VSP) protein was found in Purkinje neurons throughout the cell, which the authors suggested means Gg-VSP is expressed on the plasma membrane as well as intracellular membranes (30). Lastly, mouse VSP (Mm-VSP) was found in the brains of adult and neonatal mice, specifically in dissociated cortical neurons (32). However, to our knowledge, a connection between VSP activity and PIP-mediated biological processes in native tissues remains unknown.

To better understand VSP’s biological role, we tested for Xl-VSP localization in *Xenopus laevis* embryos. *X*. *laevis* are an allotetraploid species and have two highly similar VSP proteins, termed Xl-VSP1 and Xl-VSP2 (22,33). We found the RNA transcripts for both in multiple stages of embryonic development. The transcript expression patterns differed between the two homologs, suggesting that they fulfill different biological roles. We validated a new VSP antibody, N432/21 (NeuroMab), which recognizes both Xl-VSPs and shows that Xl-VSP protein is expressed in the brain and kidney of adult and embryonic *X*. *laevis*. Furthermore, we found that Xl-VSP protein is specifically located on the apical membrane of both embryonic and adult kidney tubules. To probe the function of Xl-VSP in these tubules, we tested the catalytic activity of both Xl-VSP homologs. We found that they dephosphorylate the 5-phosphate from PIPs, but unlike other VSPs, Xl-VSP2 is a significantly weaker 3-phosphatase. From our results, we suggest that by dephosphorylating PIPs in a voltage-dependent manner, Xl-VSPs play a fundamental role in kidney development and function.

## Results

### Xl-VSP mRNA transcripts in *X. laevis* embryos

The location and function of a biological molecule are often tightly coupled characteristics. Our previous publication showed adult *X*. *laevis* transcript expression in several tissues including testes, kidney, ovary, liver, and brain (22). Although previous studies found VSP transcripts in the embryos of ascidians (28), fish (31), chicken (21,30) and mice (27), expression in *X. laevis* embryos has not been reported. To better understand Xl-VSP’s physiological role, we determined the RNA transcript expression of Xl-VSP1 and 2 during *X*. *laevis* development. We made polyadenylated cDNA libraries from the total RNA from Nieuwkoop and Faber (NF) (34) stage 12-40 embryos and conducted semi-quantitative reverse transcriptase PCR (sqRT-PCR) using Xl-VSP1 and Xl-VSP2 specific PCR primers. We found that both Xl-VSP1 and 2 transcripts are expressed in the developing embryo and accumulate by swimming tadpole stages (NF stages >36; Fig 1A-B). Furthermore, we observed a high level of Xl-VSP1 transcripts in early embryos, which are likely remnants of maternal transcripts in agreement with our previous finding of high level expression of Xl-VSP1 in the ovary (Fig 1A) (22). Our results indicate that both Xl-VSP1 and Xl-VSP2 transcripts are present at early stages of *X. laevis* development and that the two show different patterns of embryonic RNA expression. These results agree with our previous finding that the transcripts are differentially expressed in adult tissues (22) and further suggest discrete functional roles for each in developing embryos.

**Fig 1:**
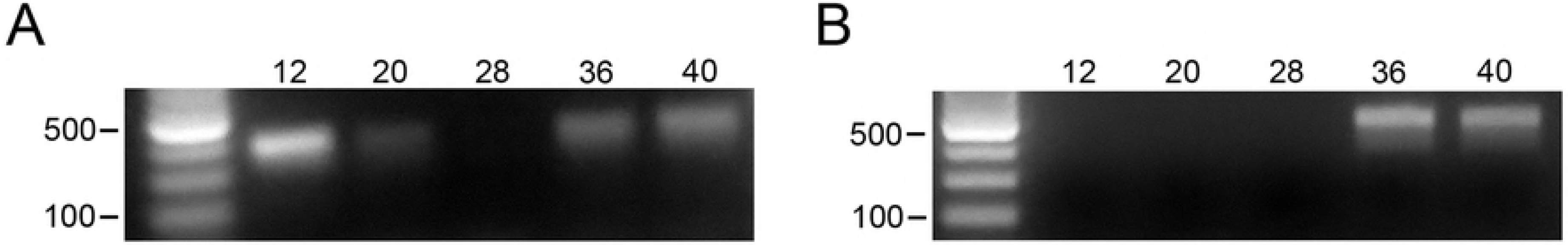
*X. laevis* embryonic stages show VSP mRNA transcripts expression. (A-B) Semi-quantitative RT-PCR (sqRT-PCR) of a panel of *X. laevis* embryos (NF stage 12-40) using PCR primers specific for Xl-VSP1 (A) and Xl-VSP2 (B). Both Xl-VSP1 and Xl-VSP2 transcripts appear to accumulate by stage 36. No bands were seen without reverse transcriptase (not shown). All cDNA libraries were made with equal amounts of total RNA as determined by spectrophotometry and confirmed by agarose gel electrophoresis to visualize ribosomal RNA bands (S1 Fig). sqRT-PCR was repeated at least two times with at least two different embryonic cohorts. The expected PCR amplicon sizes are 389 bp for Xl-VSP1 and 478 bp for Xl-VSP2. Shown are representative gels.

We next localized Xl-VSP mRNA in embryos by *in situ* hybridization (ISH). We tested NF stage 32-36 *X. laevis* embryos by ISH with an antisense-stand probe against full-length Xl-VSP1 mRNA since our sqRT-PCR analysis showed accumulation of mRNA expression at those stages. In agreement with our sqRT-PCR results, we found RNA transcripts in the developing pronephros, brain and brachial arches (Fig 2A), whereas no staining was seen when using a sense-strand control probe (Fig 2B). Since Xl-VSP1 and Xl-VSP2 mRNAs share 93% identity on the nucleotide level, it is possible this staining is due to transcripts from either Xl-VSP1, Xl-VSP2, or both.

**Fig 2:**
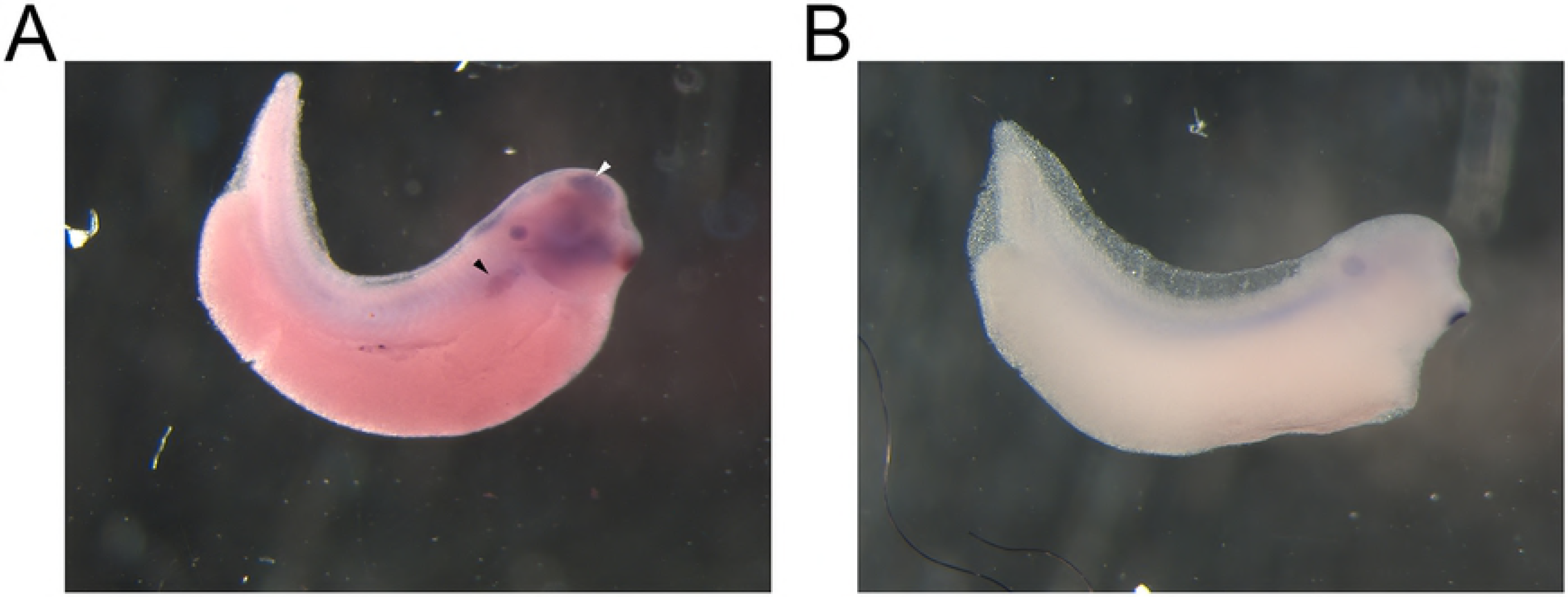
VSP mRNA is located in the pronephros and brain of *X. laevis* embryos. (A) *In situ* hybridization (ISH) of whole-mount NF stage 32 embryos. An anti-sense probe against Xl-VSPs shows Xl-VSP transcript in the proximal pronephritic field (black arrowhead) and brain (white arrowhead) of the embryos. This probe cannot distinguish between Xl-VSP1 and Xl-VSP2 mRNAs because of the similarity between the two transcripts at the nucleotide level (93%). (B) No staining was observed by ISH in a sibling embryo with a sense control probe. ISH was repeated four times with four different embryonic cohorts. Shown are representative embryos.

### Validating a VSP antibody

To determine Xl-VSP protein expression in tissues, we validated a mouse monoclonal VSP antibody, N432/21, developed by the NeuroMab facility at the University of California, Davis. We used *X laevis* oocytes as a heterologous expression system to determine the specificity of the anti-VSP. Oocytes were injected with 20 ng of VSP RNAs from different species, incubated for 36 hours before being lysed, and the lysates were run on SDS-PAGE gels for Western blotting. The anti-VSP N432/21 recognized several species of VSP, including Xl-VSP1 (58 kDa), Xl-VSP2 (58 kDa), *Xenopus tropicalis* VSP (Xt-VSP, 58 kDa), FLAG-tagged *Ciona intestinalis* VSP (Ci-VSP, 66 kDa), and *Danio rerio* VSP (Dr-VSP, 58 kDa) (Fig 3A). Xl-VSPs heterologously-expressed in *X. laevis* oocytes often displayed a double band at their predicted weight (58 kD), with the Xl-VSP2 migrating slightly slower through the gel than Xl-VSP1 and Xt-VSP (Fig 3A-C). The nature of these doublets and the slight difference in electrophoretic mobility between Xl-VSP1, Xl-VSP2, Xt-VSP, and Dr-VSP (despite their nearly identical predicted MWs) is not presently understood. These results demonstrate that anti-VSP N432/21 recognizes both Xl-VSP1 and 2 proteins, as well as the most commonly-studied VSP (Ci-VSP) and VSPs from two other common vertebrate models of development.

**Fig 3:**
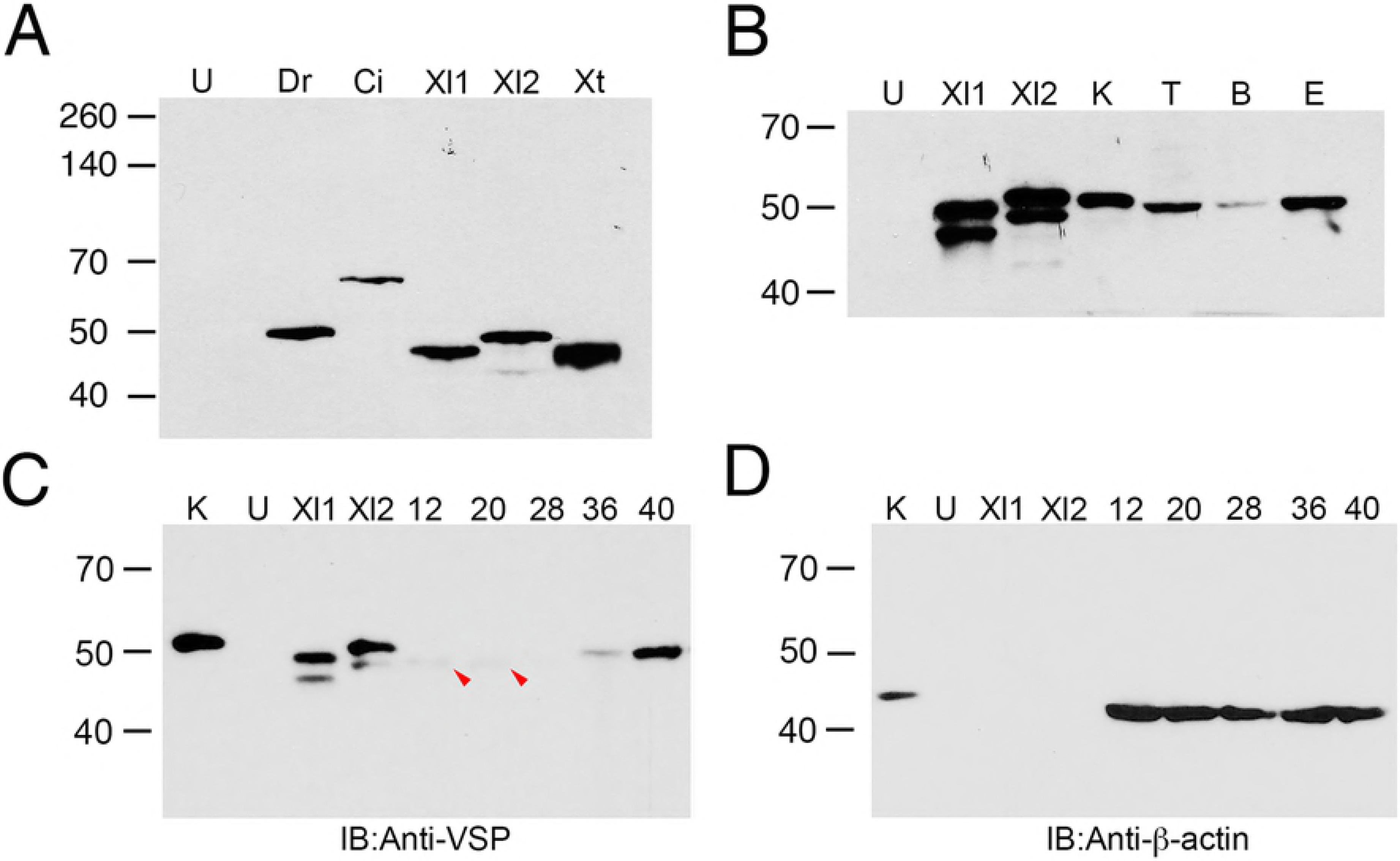
*X. laevis* tissues and embryos show VSP protein expression. (A) Western blot validation of N432/21 anti-VSP in *X. laevis* oocytes injected with RNA for Dr-VSP (Dr), FLAG-Ci-VSP (Ci), Xl-VSP1 (Xl1), Xl-VSP2 (Xl2), Xt-VSP (Xt), or left un-injected (U). All VSPs tested were recognized by the antibody. Un-injected oocytes (U) display no band. Dr-VSP, Xl-VSP1, Xl-VSP2 and Xt-VSP have a predicted MW of 58 kD while FLAG-Ci-VSP has a predicted MW of 66 kD. The slight difference in electrophoretic mobility between VSPs and their predicted MWs and the nature of the double band for Xl-VSP1 and Xl-VSP2 (as seen in panels B and C) has not been determined. These results show that anti-VSP N432/21 is specific for VSP and cross-reacts with VSPs from multiple species. (B) Western blot analysis of *X. laevis* tissues. Lysates from adult kidney (K, 3 μg), testis (T, 30 μg), brain (B, 10 μg), and stage 44 embryo (E, 20 μg) were run against lysates from oocytes injected with RNAs for Xl-VSP1 (Xl1), Xl-VSP2 (Xl2) or left un-injected (U) and analyzed by Western blot with anti-VSP. A single band of approximately the correct MW (58 kD) was observed in the tissue lysates, indicating the presence of Xl-VSP protein in all tissues tested. (C-D) Western blot analysis of *X. laevis* embryos. Lysates from NF stage 12-40 embryos (30 μg each) were run against lysates from adult kidney (K, 5 μg) and oocytes injected with RNAs for Xl-VSP1 (Xl1), Xl-VSP2 (Xl2) or left un-injected (U). Blots were probed either with anti-VSP (C) or anti-actin (D) as a loading control (predicted MW 42 kD). A weak band (potentially corresponding to Xl-VSP1) is present only at early embryonic stages 12-20 (red arrowheads), whereas a slightly slower-migrating band (potentially corresponding to Xl-VSP2) accumulates at later embryonic stages 36-40. Lysates, gels and blots were repeated three times with either three different adults or three different embryonic cohorts. Shown are representative gels for each.

### Western blots of Xl-VSP protein in embryos and adult tissues

After validating the VSP antibody, we turned to tissue samples from adult and embryonic *X. laevis*. Lysates made from adult kidney, testis, brain, and NF stage 44 embryos show a single band of the correct approximate MW (58 kD) by Western blot, indicating that these tissues express Xl-VSPs (Fig 3B). The anti-VSP N432/21 recognizes both Xl-VSP1 and 2 so we cannot definitively distinguish whether Xl-VSP1, Xl-VSP2, or both are present. However, their slight difference in electrophoretic mobility makes it possible to infer that Xl-VSP2 is predominantly expressed in the kidney while Xl-VSP1 expression may predominate in the testes, further supporting the likelihood of different roles for the two Xl-VSP proteins.

Next, we tested for Xl-VSP protein expression in *X. laevis* embryos, NF stages 12-40. Western blots show a faint band at the early embryonic stages, NF 12-20, and darker bands at the later swimming tadpole stages, NF 36-40 (Fig 3C). Based on the slight electrophoretic differences observed in the heterologous system, the slightly lower apparent MW faint bands may represent Xl-VSP1. The darker bands have a slightly higher apparent MW, suggesting they represent Xl-VSP2. Interestingly, these results suggest that Xl-VSP2 is up-regulated as the tadpole develops. This pattern of Xl-VSP protein levels agrees with the high level of Xl-VSP1 maternal transcripts in the ovary (22) and up-regulation of Xl-VSP2 RNA during later developmental stages (Fig 1B). Furthermore, from these blots, we suggest post-transcriptional regulation of Xl-VSP protein expression in the developing embryo since Xl-VSP1 transcripts also are present in later embryonic stages yet are not detectable by Western blot (Fig 3C). Overall, Xl-VSP protein is present in both adult and embryonic *X. laevis.* The differential protein expression between Xl-VSP1 and Xl-VSP2 is consistent with the differences observed at the RNA level and further suggests the two homologs serve different functional roles.

### Immunohistochemistry of Xl-VSP proteins in kidney and brain

Since protein accumulation was not detected by immunoblotting until later embryonic stages, we next used whole-mount indirect IHC on NF stage 42 embryos. Because we observed high RNA and protein levels in the kidneys, we initially focused on kidney expression for the IHC experiments. To facilitate kidney identification, we used two transgenic lines that express GFP in the developing kidney, either Pax8:GFP (35) or Cdh17:GFP (36). Whole-mount immunostained Pax8:GFP embryos showed distinct Xl-VSP staining in the proximal pronephros, indicating Xl-VSP protein is present during kidney development (S2 Fig A-C, A’-C’). This staining was not seen in sibling embryos stained with the secondary antibody alone (S2 Fig A’’-C’’). Focusing on the pronephritic tubules, we fixed, embedded, and sectioned NF stage 42 Cdh17:GFP embryos for IHC to determine subcellular localization of Xl-VSPs (Fig 4A-F). Interestingly, we observed Xl-VSP staining in the developing pronephritic tubules, specifically localized to the lumenal surface or apical membrane of the tubules (Fig 4A). This is in contrast to the soluble GFP marker that is visible throughout the tubules (Fig 4B). Consecutive sections probed with the secondary antibody alone show the GFP marker without any Xl-VSP staining (Fig 4A’-F’). We also examined embryos for Xl-VSP protein in brain tissue, given the presence of Xl-VSP RNA transcripts in embryonic brain (Fig 2A). However, we did not observe any Xl-VSP staining in the developing brain, either in whole-mount or in sections (data not shown). This may be due to Xl-VSP protein levels being too low to visualize via our IHC methods, or due to masking of the epitope during fixation by Xl-VSP binding partners (see discussion). Further studies involving IHC signal amplification and/or antigen retrieval may be needed to distinguish between these possibilities and to localize Xl-VSP protein in the *X. laevis* brain. Our findings demonstrate embryonic expression of Xl-VSP protein in pronephritic tubules with subcellular localization on their apical membranes, thus, suggesting Xl-VSPs play an important role in the development and function of the embryonic *X. laevis* kidney.

**Fig 4:**
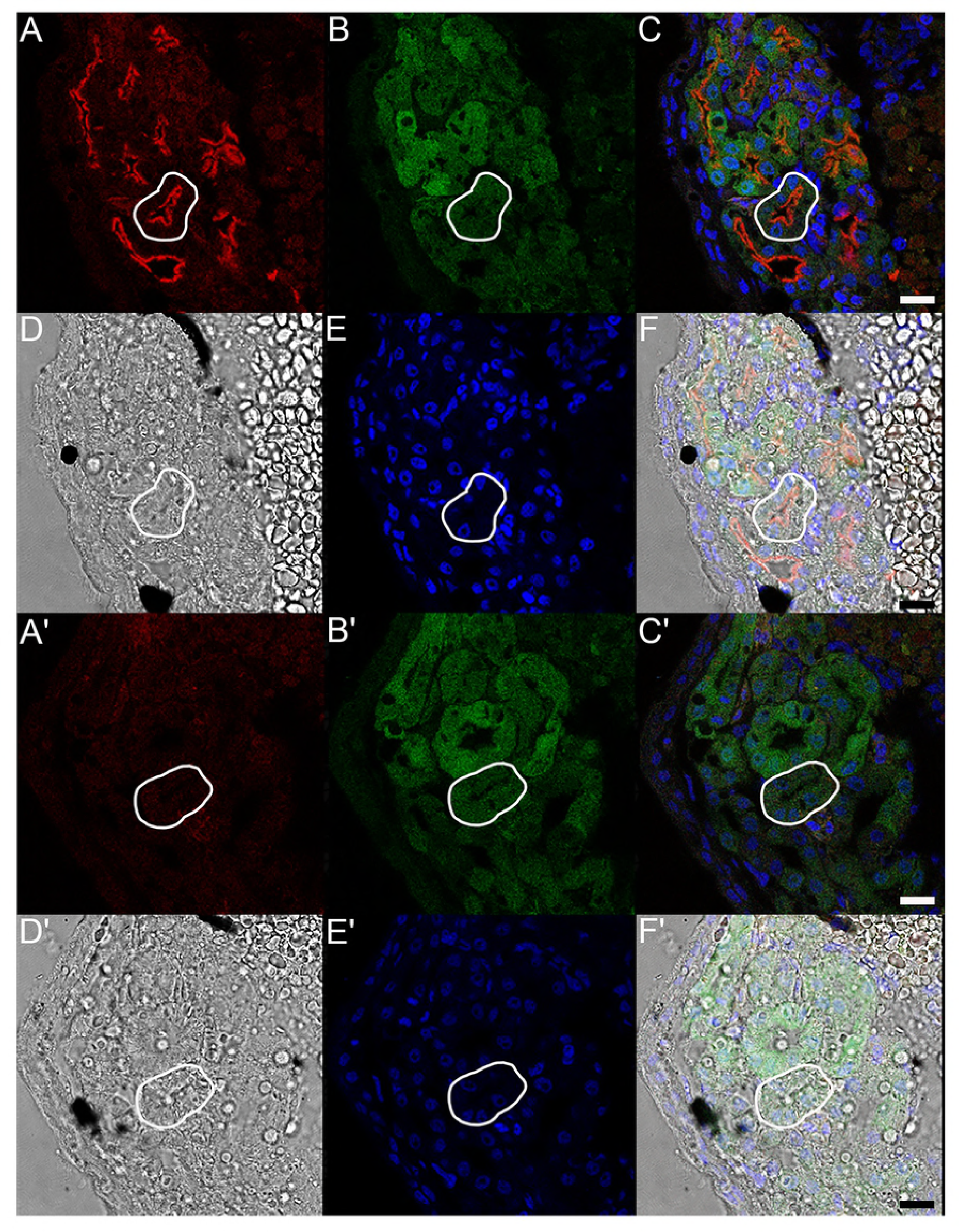
Xl-VSP protein is expressed on the lumenal surface of embryonic pronephroi. NF stage 42 Cdh17:GFP *X. laevis* embryos were sectioned and stained to test for VSP protein. Kidney tubules were identified by the presence of the Cdh17:GFP transgene (B, B’). Sections stained with anti-VSP (A) showed fluorescence on the lumenal surface (corresponding to the apical membrane) of the proximal kidney tubule cells (outlined in white). Anti-VSP staining was only observed in the presence of anti-VSP (A-F) and not in secondary antibody alone control sections (A’-F’). Panels (A, A’) anti-VSP or no anti-VSP control; (B, B’) GFP transgene; (C, C’) fluorescence signal overlay of A, B and E; (D, D’) bright field images; (E, E’) Hoechst 33342 (to mark nuclei); (F, F’) signal overlay of A, B, D and E. IHC was repeated three times with three different embryonic cohorts. Shown are representative sections. Scale bar = 25 μm

In addition to embryos, we also tested adult kidney tissue using IHC. Adult male *X*. *laevis* kidneys were removed, fixed, embedded, sectioned, and stained with anti-VSP. The sections also show Xl-VSP staining on the lumenal surface of the tubules (Fig 5A), which is consistent with the subcellular staining observed in embryonic kidney tubules (Fig 4A). No staining was seen in adult kidney tubules when consecutive sections were probed with secondary antibody alone (Fig 5B). In contrast to our IHC of embryonic kidney sections, we only observed staining in adult kidney sections by anti-VSP after post-fixation antigen retrieval, which suggests a difference in the availability of the VSP antigen in embryonic versus adult kidney tissues. These results suggest a long-term functional role for Xl-VSPs in the kidney since they are present from the earliest stages of kidney development and continue into adulthood.

**Fig 5:**
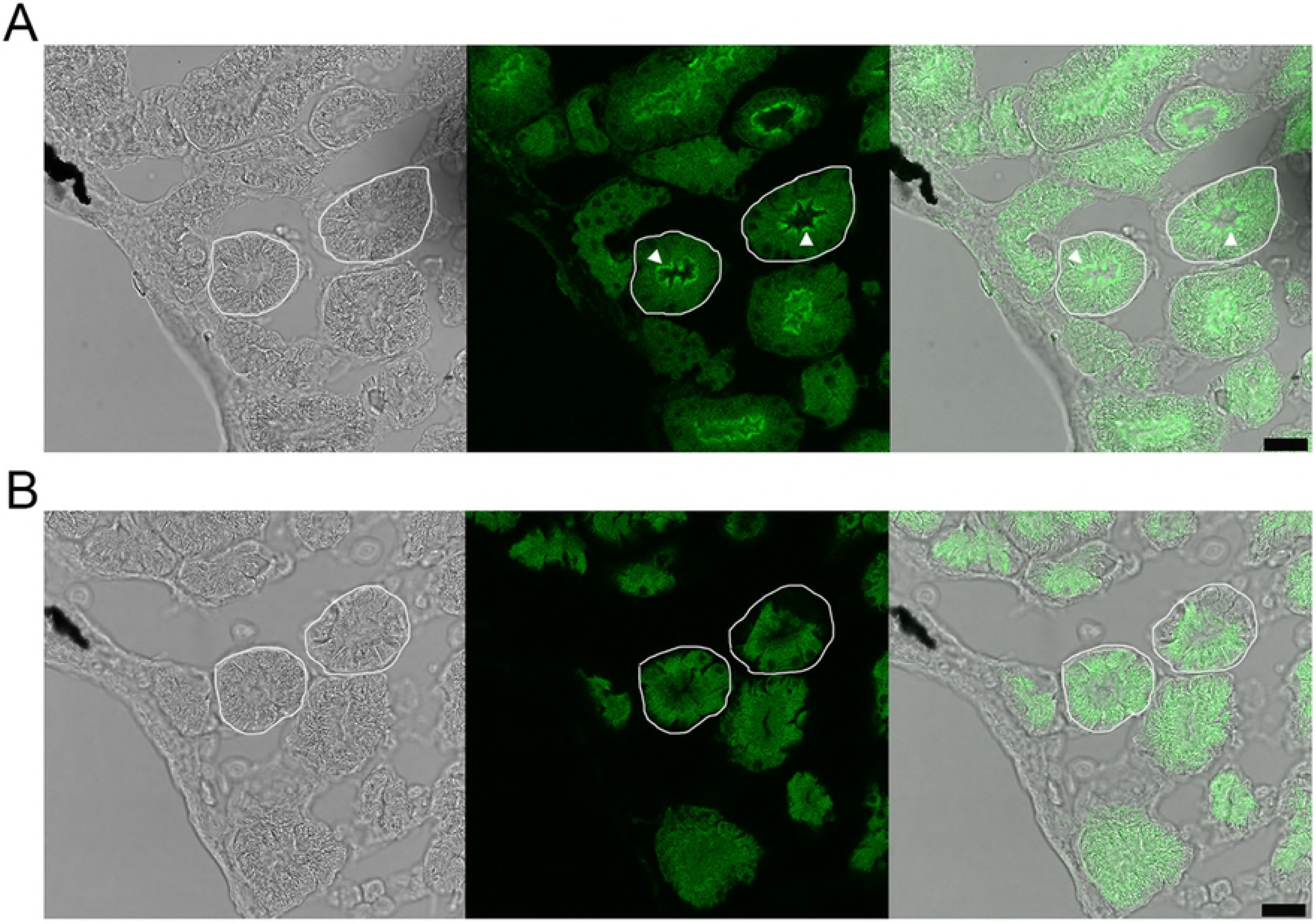
Xl-VSP protein is expressed on the lumenal surface of adult kidney tubule epithelial cells. (A) Adult kidney sections were stained with anti-VSP and anti-mouse IgG-Alexa 488. Xl-VSP staining is observed on the lumenal surface (marked with arrowheads) of the kidney tubules (two outlined in white). This surface corresponds to the apical membrane and not the basolateral membrane of the epithelial cells. (B) Apical membrane staining was not seen on a consecutive section in the absence of anti-VSP primary antibody. IHC was repeated on sections from kidneys of three different adult males. Shown are representative sections. Scale bar = 25 μm

### Functional activity of Xl-VSPs

While VSPs from *Ciona intestinalis* and *Danio rerio* function as both 5- and 3-phosphatases (10–12,14,37) the Xl-VSPs have previously been characterized as only a 5-phosphatase against phosphatidylinositol-4,5-bisphosphate [PI(4,5)P_2_] (22). To understand how the Xl-VSPs may contribute to kidney function, we examined their phosphatase activity to determine whether both Xl-VSP1 and 2 dephosphorylate phosphatidylinositol-3,4,5-trisphosphate [PI(3,4,5)P_3_] at the 5-phosphate as well as whether they function as 3-phosphatases. Because Xl-VSPs are regulated by voltage, we used two electrode voltage clamp (TEVC) to control Xl-VSP activation. To monitor all four possible dephosphorylation reactions (Fig 6A), we turned to pleckstrin homology (PH) domain-based optical biosensors. PH domains are well known proteins that bind specifically to certain PIPs (38-40). We chose PH domains from tandem PH domain containing protein 1 (TAPP) and phospholipase C (PLC) because they bind specifically to phosphatidylinositol-3,4-bisphosphate [PI(3,4)P_2_] and PI(4,5)P_2_, respectively, allowing us to monitor both 5- and 3-phosphatase activities (12,14,41,42). To follow PI(3,4)P_2_, we used the biosensor fTAPP that utilizes the PH domain from TAPP flanked by an N-terminal CFP and C-terminal YFP, while the whole sensor is anchored to the membrane through a prenylation site at the C-terminus (Fig 6B) (41-43). When VSP dephosphorylates the 5-phosphate from PI(3,4,5)P_3_, the TAPP-PH domain of the fTAPP biosensor will bind the PI(3,4)P_2_ and the resulting conformational change between the CFP and YFP will change the Förster resonance energy transfer (FRET) between them, leading to an increase in the FRET ratio (Fig 6B). As PI(3,4)P_2_ is depleted when VSP dephosphorylates the 3-phosphate, the TAPP-PH unbinds and the FRET ratio decreases (Fig 6B). Thus, we combine the optical measurements of the biosensors with TEVC to precisely control and monitor catalytic activity of Xl-VSPs.

**Fig 6:**
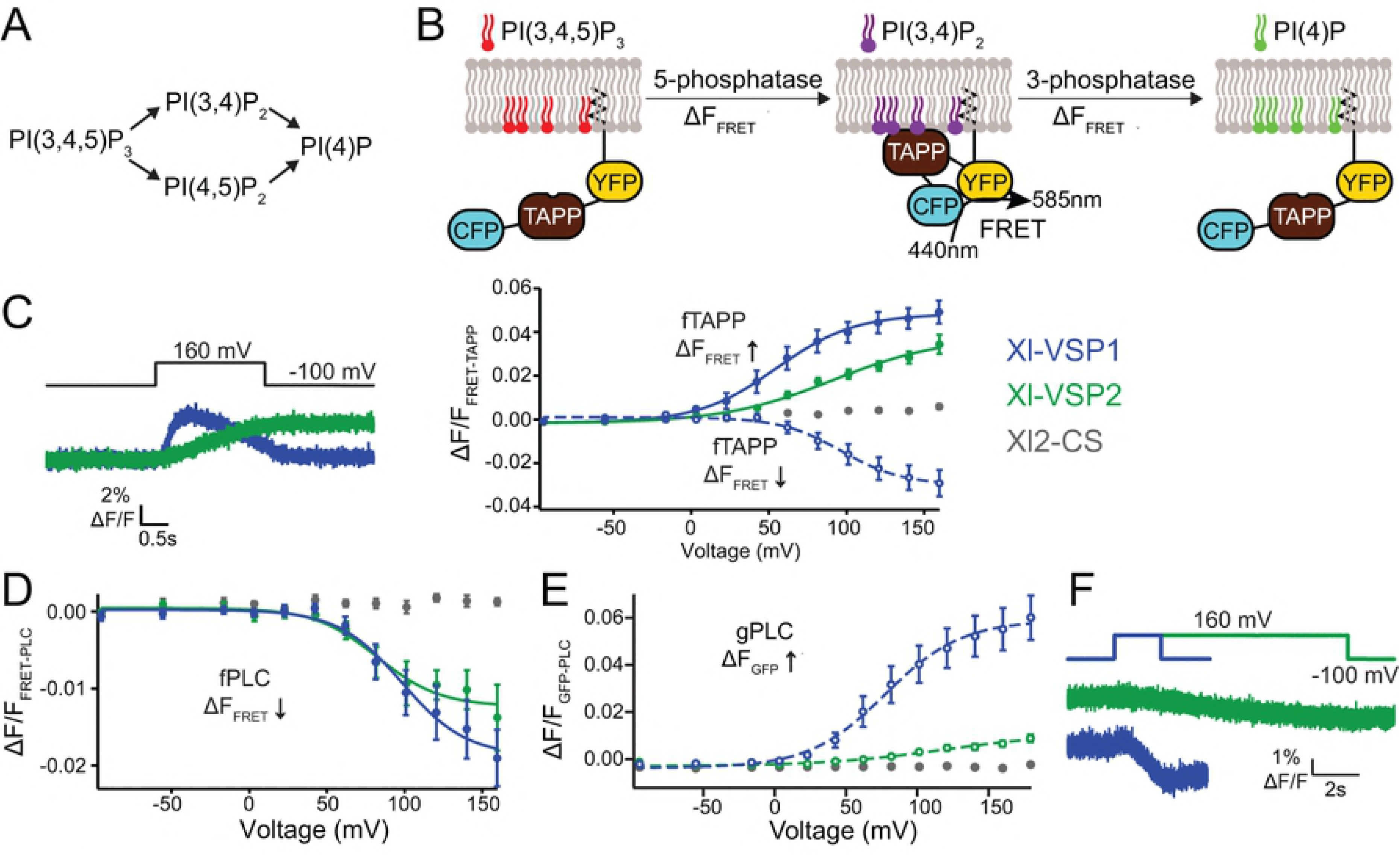
Xl-VSP1 and Xl-VSP2 function as voltage-regulated 5- and 3-phosphatases. (A) Schematics of the known VSP reactions. (B) Cartoon representation of fTAPP binding and release with increasing and decreasing PI(3,4)P_2_ concentrations. The binding of fTAPP to PI(3,4)P_2_ results in a conformational change that increases the FRET signal. Similarly, a reduction of PI(3,4)P_2_ results in a decrease in the FRET signal. (C) Oocytes were injected with fTAPP and either Xl-VSP1, Xl-VSP2 or Xl-VSP2 C301S (Xl2-CS), a catalytically inactivated protein. (left) Averaged fTAPP FRET traces over time during a voltage step from a holding potential of -100 to +160 mV. The FRET signal increases and decreases in the same pulse for Xl-VSP1 while it only increases for Xl-VSP2. (right) FRET measurements were tested at several voltages and the ΔF/F fTAPP FRET ratio was plotted versus the voltage (RV). The FRET increase (net 5-phosphatase reaction, solid line, closed symbols) was plotted separately from the FRET decrease (net 3-phosphatase reaction, dashed line, open symbols). Xl-VSP1 shows robust activity as both a 3- and 5-phosphatase while Xl-VSP2 functions as a 5-phosphatase. (D) Oocytes were injected with fPLC and either Xl-VSP1, Xl-VSP2 or Xl2-CS. The ΔF/F fPLC FRET RV shows a FRET decrease (net 5-phosphatase reaction, solid line, closed symbols) for both Xl-VSP1 and 2 indicating both dephosphorylate PI(4,5)P_2_. (E) Oocytes were injected with gPLC and either Xl-VSP1, Xl-VSP2 or Xl2-CS. GFP fluorescence measurements were tested at several voltages and the fluorescence voltage relationship plotted showing a fluorescence increase (net 3-phosphatase reaction, dashed line, open symbols). While Xl-VSP1 shows significant levels of 3-phosphatase activity against PI(3,4,5)P_3_, Xl-VSP2 is a much less efficient 3-phosphatase. (F) Averaged fPLC FRET traces over time during a voltage step from a holding potential of -100 to +160 mV for fPLC co-expressed with either Xl-VSP1 or Xl-VSP2. The kinetics of activation are significantly faster for Xl-VSP1 than for Xl-VSP2. All error bars are ± SEM., n ≥ 8. Data fit with single Boltzmann equations

When oocytes co-expressed Xl-VSP1 and fTAPP, we observed Xl-VSP1 inducing an initial increase and subsequent decrease in FRET over time, indicating that it functions as both a 5- and a 3-phosphatase (Fig 6C, left). To determine the voltage dependence of activity, we tested several voltages and plotted the FRET versus voltage relationships. To better understand each reaction, we separated the FRET increase from the FRET decrease during our analysis (Fig 6C, right, blue). Note the voltage dependence of the Xl-VSP1 5-phosphatase reaction (FRET increase, V_1/2_ = 55 mV) is lower than the voltage dependence of the 3-phosphatase (FRET decrease, V_1/2_ = 97 mV) (S1 Table). As a control, we also tested a catalytically inactive Xl-VSP2 C301S (Xl2-CS) (22) with fTAPP. As expected, a slight, continuous FRET increase was observed. This small signal is expected because it has been previously shown that a low level of endogenous Xl-VSP activity is present in *X*. *laevis* oocytes (14,22,41,42,44). Interestingly, the same fTAPP experiment with Xl-VSP2 only showed a FRET increase (Fig 6C, green) even when we extended the voltage pulse to 60 seconds (data not shown). Additionally, as was previously reported (22), the voltage dependence of Xl-VSP2 5-phosphatase activity against PI(3,4,5)P_3_ is shifted to higher voltages (V_1/2_ = 92 mV) compared to that of Xl-VSP1 (V_1/2_ = 55 mV) (S1 Table). These results indicate that, although the Xl-VSP homologs are 93% identical, they do not share identical voltage-dependent functional roles in the cell.

To follow PI(4,5)P_2_, we used two different biosensors based on the PLC-PH domain, one FRET sensor which utilized the PH domain from PLC instead of TAPP to create fPLC, and a fluorescence sensor where GFP is attached to the N-terminus of PLC, called gPLC. We used fPLC to monitor the 5-phosphatase activity during the PI(4,5)P_2_ to PI(4)P reaction. When oocytes co-expressed either Xl-VSP1 or Xl-VSP2 with fPLC, we observed robust FRET ratio decreases (Fig 6D), indicating 5-phosphatase activity for both homologs, as was previously published (22). We used gPLC to monitor the 3-phosphatase reaction, the PI(3,4,5)P_3_ to PI(4,5)P_2_ reaction, because the gPLC sensor is a more sensitive sensor for this reaction than fPLC. Instead of a change in FRET, the gPLC translocates to the membrane when PI(4,5)P_2_ concentrations increase, resulting in an increase in fluorescence. When oocytes co-expressed Xl-VSP1 with gPLC, we observed an increase in fluorescence, indicating the production of PI(4,5)P_2_ from the Xl-VSP 3-phosphatase reaction (Fig 6E, blue). Unexpectedly, the same experiment with Xl-VSP2 also gave a small, but reproducible fluorescence increase (Fig 6E, green), indicating Xl-VSP2 is also able to function as a 3-phosphatase. However, we did not observe any 3-phosphatase activity when using the fTAPP sensor. These results show that Xl-VSP2 can remove the 3-phosphate from PI(3,4,5)P_3_ but not from PI(3,4)P_2_. Further investigations are needed to better understand this striking difference in substrate specificity. Interestingly, the kinetics of Xl-VSP2 are significantly slower than the kinetics of Xl-VSP1 (Fig 6F), as seen by the voltage steps to 160 mV for a 2 second voltage step for Xl-VSP1 and a 10 second voltage step for Xl-VSP2 (Fig 6F). Overall, our results show that Xl-VSP1 can catalyze the same four reactions as other VSPs while Xl-VSP2 is restricted by substrate. Since the reactions and kinetics appear to be different between Xl-VSP1 and 2, they likely fulfill different functional roles in tissues which correlates with their differential expression patterns.

## Discussion

VSPs provide a direct link between the electrical state of a cell and its PIP signaling pathways. By dephosphorylating PIPs in a voltage dependent manner, VSPs could regulate PIP-dependent processes such as cortical cytoskeleton remodeling, ion channel function, G protein-coupled receptor signaling (GPCR) and intracellular calcium signaling (18,19,45). Though the biophysical properties of VSPs have been the subject of intense study since the discovery of Ci-VSP (10), the physiological relevance of this direct link remains unclear.

To elucidate Xl-VSPs’ biological role, we determined the cellular expression pattern of Xl-VSP RNA and protein in *Xenopus laevis*. We extended our previous findings in adult *X*. *laevis* tissues (22) by showing both Xl-VSP1 and 2 RNA transcripts in developing embryos. Though VSP is seen in the embryonic stages of other species, this is the first study showing Xl-VSP RNA and protein in *X. laevis* embryos. Our findings indicate that Xl-VSP RNAs are present at early stages of development, NF stages 12-40, with both Xl-VSP1 and 2 RNA levels increasing from NF stage 20 to 40. At the same stages, we observed an increase in protein levels by Western blot, consistent with our RNA detection, indicating an upregulation of the protein during development. Using ISH, we observed Xl-VSP RNA transcripts in both the brain and pronephros of NF stage 32-36 embryos. Using IHC, we found Xl-VSP protein in the pronephroi of NF stage 42 embryos. Upon sectioning the embryos, we observed anti-VSP staining on the lumenal surface of the pronephritic tubules, corresponding to the apical membrane of the epithelial cells lining the tubule lumen. Additionally, we sectioned adult kidneys and found the same apical membrane localization in the tubules. To directly test Xl-VSP catalytic activation, we combined electrophysiology with optical biosensors to show that both Xl-VSPs dephosphorylate the 3- and 5-phosphate from PIPs though with different substrate specificities and efficiencies. Combining our localization results with our electrophysiological activity data, we suggest that Xl-VSPs have specific functional roles in both the brain and kidney.

Concentrating on the kidney, we found Xl-VSPs spanning the earliest stages of the developing pronephros into the full adult kidney, suggesting Xl-VSPs modulate PIP concentrations on the apical membrane of the kidney epithelial cells in a voltage dependent manner. Kidney tubules are lined with specialized epithelial cells that regulate vertebrate solute homeostasis (46,47). The transepithelial potential difference across these cells contributes to tubular reabsorption of solutes and water (48,49). The lumenal surface of these epithelial cells are decorated with microvilli and primary cilia, both of which function in tubular reabsorption by sensing fluid flow (50-54). Microvilli provide a large absorptive membrane surface for kidney epithelial cells and are actin-dependent dynamic structures with turnover rates in the tens of minutes (55). Heterologously expressed Gg-VSP influences actin-based cytoskeleton rearrangements in cultured chick fibroblasts leading to morphological changes in cortical cell structure (56), suggesting Xl-VSPs could modify the actin-based microvilli in kidney epithelial cells. In addition, GPCR signaling originating in microvilli at the apical membrane of proximal kidney tubules is directly influenced by local PI(4,5)P_2_ concentrations (57), which is a substrate for both Xl-VSPs. Likewise, primary cilia are dependent on local PI(4,5)P_2_ concentrations (58), and importantly, trigger calcium entry due to the mechanical forces from the fluid flow (54). As calcium enters the cells, the resulting depolarization could lead to activation of the Xl-VSP on the apical membrane and subsequent regulation of the cortical cytoskeleton, GPCRs, and ion channels during renal tubule development and adult renal function.

Our results are consistent with VSP homologs having a broad role in multiple tissues. Specifically, our study agrees with previous studies that also showed embryonic kidney tubule expression of VSP in zebrafish (31), and chicks (21). In particular, Gg-VSP RNA was found on the proximal part of the nephrogenic tubules of HH stage 26 chicks and not the distal or collecting tubules, consistent with our IHC results showing proximal tubule localization in NF stage 32 embryos (S2 Fig). Our findings in *X*. *laevis* are also consistent with the findings in chicks in that not all tubules appear to immunostain with our VSP antibody, indicating that the Xl-VSPs may be restricted to a functional subset of the developing and adult kidney. Our results go further because we observe clear Xl-VSP protein immunostaining on the apical membrane of the tubules where reabsorption of nutrients occurs and where PIP-mediated signaling originates.

Not all VSP studies analyzed kidney expression patterns. The human VSP, Hs-VSP1 (previously called TPIP and TPTE2), was not examined in kidneys, making human VSP kidney expression unknown (20). In addition to kidney expression, VSP is also expressed in the brain and nervous system of multiple species (20,22,28,30,32). Indeed, we observed Xl-VSP RNA transcripts and protein in the brain of adult and embryonic *X. laevis*. Some reports of VSP expression conflict, particularly regarding expression of mouse VSP, Mm-VSP (previously called PTEN2). Early studies identified Mm-VSP as a testis specific protein (25) while subsequent investigations report expression in the brain (32). While more research is needed, a pattern is emerging with the kidney and brain being the most consistent tissues expressing VSP. Our results support this pattern and a functional role for VSP in both the brain and kidney of multiple vertebrate species.

The subcellular localization of VSP is also debated in the field. Prior studies of native VSP protein have found VSP on the plasma membrane of ascidian sperm (10) and of Purkinje neurons in chick cerebellum sections (30). Along with plasma membrane localization, however, the cerebellum sections also showed intracellular staining with a Gg-VSP antibody. A more recent study in mice used dissociated cortical neurons from P0-1 mice and the staining appears throughout the cells (32). In addition, heterologous expression of mouse and human VSP consistently show intracellular membrane localizations (20,25,29). In contrast, our subcellular localization of endogenous Xl-VSP protein is on the apical membrane of kidney tubule epithelia. The differences in our results and those from prior studies may lie in the tissues tested, the antigen retrieval discussed below, or in the antibodies used. Further experiments are needed to determine whether VSPs function on both intracellular and plasma membranes. While not tested against either chicken or mouse VSPs, our antibody does cross react with VSPs from *D. rerio*, *X. tropicalis and C. intestinalis*. As a result, N432/21 may prove to be an invaluable tool in future studies of VSP’s physiological role in vertebrate kidney development and neural function.

It is interesting to note that unlike in fixed embryonic kidney tissue, we only observed staining of adult kidney tissue after post-fixation antigen retrieval. This difference in VSP epitope availability could be due to different roles for VSP in adult versus embryonic kidney tubules, and differential VSP binding partners in these tissues. We recently demonstrated that Ci-VSP forms dimers (42). Since the two Xl-VSP homologs shown here display different expression in the embryo and adult frog, it is also possible that the difference we observed in VSP epitope availability is due to the presence of Xl-VSP homodimers and heterodimers.

In addition to our localization experiments, we also tested the electrophysiological characteristics of both Xl-VSP homologs. We controlled their activation using TEVC and monitored the production and depletion of different PIPs using optical biosensors. Our previous report indicated that both homologs catalyzed the dephosphorylation of the 5-phosphate from PI(4,5)P_2_. VSPs from other species have been shown to utilize PI(3,4,5)P_3_ and PI(3,4)P_2_ as substrates as well. Here, we tested all three substrates by using PIP sensors that are sensitive to either PI(3,4)P_2_ or PI(4,5)P_2_ concentrations (TAPP-PH or PLC-PH respectively), allowing us to monitor the 3-phosphate removal from PI(3,4,5)P_3_ and PI(3,4)P_2_ as well as the 5-phosphate removal from PI(3,4,5)P_3_ and PI(4,5)P_2_. We found that Xl-VSP1 behaves much like other VSPs in that it dephosphorylates both 5- and 3-phosphates. Xl-VSP2, on the other hand, is a slower 5-phosphatase and a weaker 3-phosphatase. After modifying our protocols for longer depolarizations to account for the slower kinetics, we observed clear 5-phosphatase activity for Xl-VSP2 using both fTAPP and fPLC sensors. The voltage-dependence of activation was shifted to higher voltages for Xl-VSP2 compared to Xl-VSP1. We also observed 3-phosphatase activity of Xl-VSP2 against PI(3,4,5)P_3_, but not against PI(3,4)P_2_, even after a one minute depolarization. This difference between the Xl-VSP homologs further indicates that the two are not interchangeable and serve different physiological roles in the cell.

In conclusion, we observe Xl-VSP expression in the brain and kidney of *X*. *laevis* embryos and adults. We localize Xl-VSP protein on the apical membrane of kidney tubule epithelial cells, and we characterize both Xl-VSPs as 3- and 5-phosphatases with different enzymatic activities. Our results indicate a role for voltage-dependent dephosphorylation of both 5- and 3-phosphates from PIPs on the apical membrane during kidney tubule development and renal function.

## Materials and methods

### Materials

*Ciona intestinalis* VSP (Ci-VSP, NM_001033826) and *Danio rerio* VSP (Dr-VSP, NM_001025458) plasmids were kindly provided by Y. Okamura (Osaka University, Osaka, Japan). *X. laevis* VSP1 (Xl-VSP1, NM_001096603), *X. laevis* VSP2 (Xl-VSP2, NM_001280607), and *X. tropicalis* VSP (Xt-VSP, NM_001015951) expression plasmids were made in Laurinda Jaffe’s laboratory and are available through Addgene (Cambridge, MA). While the original human gene was named transmembrane phosphatase with tensin homology (TPTE), the majority of the literature refer to this protein as the voltage sensing phosphatase or VSP. Here, we follow this more common nomenclature to avoid confusion. fPLC and fTAPP were both kindly provided by E.Y. Isacoff (University of California, Berkeley). GFP-PLC-PH was kindly provided by T. Meyer (Stanford University). Mutations and epitope tags were created by site-directed mutagenesis with Pfu Turbo DNA polymerase (Agilent, Santa Clara, CA) by standard protocols. All DNA constructs were confirmed by sequencing. cRNA was transcribed using SP6 or T7 mMessage mMachine kits (Thermo-Fisher, Waltham, MA). *X*. *laevis* oocytes for Xl-VSP activity assays were purchased from Xenopus 1 (Dexter, MI). Mouse monoclonal anti-VSP (N432/21; RRID:AB_2716253) was made by the UC Davis/NIH NeuroMab Facility and is available through Antibodies Incorporated (Davis, CA). Anti-beta actin (clone C4, catalog # sc-47778) was purchased from Santa Cruz Biotechnology (Dallas, TX). 3G8.2C11 antibody was obtained from the European Xenopus Resource Centre (59,60). Goat anti-mouse IgG, light chain specific, conjugated to horseradish peroxidase (HRP, catalog # 115-035-174) was purchased from Jackson ImmunoResearch (West Grove, PA). Goat anti-mouse IgG conjugated to Alexa Fluor 488 or Alexa Fluor 594 was purchased from Thermo Fisher. Anti-digoxygenin (DIG) conjugated to alkaline phosphatase (AP) was purchased from Sigma-Aldrich (St. Louis, MO). Cdh17:GFP and Pax8:GFP transgenic *Xenopus laevis* frogs were obtained through the National Xenopus Resource (Woods Hole, MA, RRID:SCR_013731) (35,36,60%). All animals in this study were used in accordance with protocols approved by the Montana State University IACUC.

### Reverse-transcriptase PCR

Total RNA was isolated from embryos with TRIzol (Thermo Fisher), quantitated by Nanodrop spectrophotometry and analyzed by ethidium bromide agarose gel electrophoresis (S1 Fig). Two μg of embryo RNA was treated with DNAse I (DNA-free, Thermo Fisher) to remove genomic DNA and cDNAs were reverse-transcribed by priming with oligo-dT using SuperScript IV reverse transcriptase (Thermo Fisher). A parallel control reaction was performed without reverse-transcriptase for each DNAse I-treated RNA. Semi-quantitative PCR was performed on 1/20^th^ of the resulting cDNA using primers for Xl-VSP1 (forward: GATGCTGGAAACAACTCCATAGTCC; reverse: GCTGTGTATGTGGTCAGAACAC) or Xl-VSP2 (forward: GATGCTGGGAACAATTCCGTAGTCA; reverse: GGGTAATAGTACGTTAAGAAGTG) with Taq polymerase in Standard Taq PCR buffer (New England Biolabs, Ipswich, MA) with the thermocycling parameters: 95°C for 30 sec, 37 cycles of 62°C for 20 seconds followed by 68°C for 30 seconds. For each primer pair the forward primer is located upstream of an intron and the reverse primer is located in the 3’ UTR to ensure alloallele specificity and amplification of only cDNA. PCR products were analyzed by ethidium bromide agarose gel electrophoresis.

### *In-situ* hybridization (ISH)

Full-length antisense and sense digoxygenin (DIG)-labeled probes were made from linearized pcDNA3-Xl-VSP1 plasmid with SP6 or T7 RNA polymerase and DIG NTPs (Sigma-Aldrich). *In situ* hybridization to detect Xl-VSP mRNAs was performed on NF stage 32-36 embryos fixed for 90 minutes in phosphate-buffered 4% paraformaldehyde (pH 7.4) by standard methods (61). Embryos were probed with anti-DIG IgG-AP, stained with NBT-BCIP (Sigma-Aldrich) and imaged with a ProgRes C14plus camera (Jenoptic, Germany) through a 5x objective on an Axioscope.A1 microscope (Zeiss, Germany).

### Defolliculating *X*. *laevis* oocytes

*X.laevis* ovaries were purchased from Xenopus 1 (Dexter, MI). Ovaries were processes as described previously (62). Briefly, each ovary was washed once and morselized in Ca^2+^-free (96 mM NaCl, 2 mM KCl, 1 mM MgCl_2_, and 10 mM HEPES, pH 7.6). The morselized ovary was washed in Ca^2+^- free to remove yolk from lysed oocytes and then digested at room temperature for 30-45 minutes in 2% collagenase (Sigma-Aldrich, catalog # C-0130), 0.1% soybean trypsin inhibitor (Sigma-Aldrich, catalog # T-9003), and 0.1% BSA (Sigma-Aldrich, catalog # A3294) made in Ca^2+^-free. Post-digestion, the oocytes were washed in oocyte wash buffer (34 mM KH_2_PO_4_, 66 mM K_2_HPO_4_, 0.1% BSA, pH 6.5) at least 10 times to remove follicles, followed by 10 washes with Ca^2+^-free. The oocytes were then sorted into ND-96 (96 mM NaCl, 2 mM KCl, 1.8 mM CaCl_2_, 1 mM MgCl_2_, 50 mg/ml gentamicin, 2.5 mM sodium pyruvate, and 10 mM HEPES, pH 7.6) and cultured at 18°C.

### Western Blotting

Adult *Xenopus laevis* tissue, oocyte, and embryo lysates were made by homogenization and brief sonication in lysis buffer (150 mM NaCl, 150 mM dithiothreitol, 0.1% CHAPS, 50 mM Tris, pH 7.4) with protease inhibitors (Roche, Switzerland). *X. laevis* oocytes were injected with 20 ng VSP RNA and cultured at 18°C for 36 hours before lysis. Protein concentrations were determined by a Quick Start Bradford assay (Bio-Rad, Hercules, CA). Lysates were run on 10% polyacrylamide gels at 200 volts for one hour, transferred to ProTran nitrocellulose membranes (GE Healthcare, Chicago, IL) in sodium borate transfer buffer for 70 minutes at 350 mA and stained with Ponceau-S to evaluate equal loading of protein between the wells. Membranes were blocked in TBS-T block buffer (5% non-fat dry milk, 150 mM NaCl, 10 mM, 0.1% Tween-80, 10 mM Tris, pH 7.4) and probed with anti-VSP supernatant at 1:25 dilution or anti-beta actin at 1:500 dilution. Blots were then washed in TBS-T, probed with anti-mouse IgG-HRP and developed with ECL reagent (Sycamore Life Sciences, Houston, TX).

### Immunohistochemistry (IHC)

Adult *Xenopus laevis* kidneys and NF stage 40-42 embryos (34) were fixed in phosphate-buffered 4% paraformaldehyde (pH 7.4) for 90 minutes followed by sequential equilibration for two hours each in 30% sucrose, 30% sucrose/50% O.C.T (Optimal Cutting Temperature) embedding media (Fisher, Hampton, NH), and 100% OCT. Kidney tissues were then snap-frozen in OCT using a dry ice/ethanol slurry and sectioned at 14 μm onto glass slides coated with 0.5% porcine gelatin, type A (Sigma-Aldrich, catalog # G2500) and 0.05% chromium potassium sulfate. Sections were air-dried overnight at room-temperature and stored at -80°C. Sections were rehydrated with phosphate-buffered saline (PBS; 137 mM NaCl, 27 mM KCl, 100 mM Na_2_HPO_4_, 18 mM KH_2_PO_4_, pH 7.4) and subjected to antigen retrieval in 10 mM sodium citrate, 0.05 % Tween-80, pH 6.0 for 20 minutes at 95°C followed by quick cooling on ice. Sections were re-equilibrated with PBS-T (0.1% Triton X-100) and blocked in PBS-T with 10% normal goat serum (Fisher, catalog # 16-210-064). Sections were probed with anti-VSP supernatant at 1:25 dilution, washed in PBS-T, and probed with anti-mouse IgG-Alexa Fluor 488 or anti-mouse IgG-Alexa Fluor 594 at 1:1000 dilution with 1 μg/mL Hoechst 33342 (Thermo-Fisher) to stain nuclei. Immuno-stained sections were mounted in Vectashield (Vector Labs, UK) under glass coverslips and imaged with a 63x/1.4 NA objective on a Leica SP8 confocal microscope (Leica, Buffalo Grove, IL). Embryo sections were similarly fixed, sectioned, stained and imaged without the antigen retrieval process. Whole-mount embryos were similarly fixed and stained without the antigen retrieval process and with four hours to overnight incubations to allow for antibody penetration. Whole-mount IHC images were captured using Progres Mac CapturePro software on a ProgRes C14plus camera through a Zeiss Plan-neofluar 5x/0.16 NA objective on an Axioscope.A1 microscope. GFP or Alexa Fluor 594 fluorescence in whole-mount immuno-stained embryos was excited by an LED and collected through Zeiss FITC (EX BP 475/40, BS FT 500, EM BP 530/50) or Texas Red (EX BP 560/40, BS FT 585, EM BP 630/75) filter sets, respectively. Anti-VSP, anti-3G8 and matching secondary-alone control samples were imaged using the same excitation intensity and digital gain. IHC was repeated a total of three times with tissue from different animals or clutches of embryos.

### Electrophysiology and fluorescence measurement of activity

Two electrode voltage clamp (TEVC) was performed as previously described (14). FRET-based PIP sensors (41,42) were used to measure depletion of PI(4,5)P_2_ or the production and depletion of PI(3,4)P_2_. The PIP sensors were designed by adding an N-terminal CFP and a C-terminal YFP to pleckstrin homology (PH) domains. The PH domains were originally derived from phospholipase C (PLC) for PI(4,5)P_2_ and the tandem PH domain containing protein 1 (TAPP) for PI(3,4)P_2_, and hence were called fPLC and fTAPP respectively. To measure production of PI(4,5)P_2_, the diffusion based sensor GFP-PLC (gPLC) was used which has a GFP was attached to the N-terminus of the PLC-PH domain. All Xl-VSP cRNAs (0.8 μg/μl) were mixed with either fPLC cRNA (0.4 μg/μl), fTAPP cRNA (0.4 μg/μl) or gPLC cRNA (0.06 μg/μl) and injected into *X. laevis* oocytes. In all experiments, 50 nl of the cRNA mixtures were injected into oocytes and incubated in ND-96 for 30-40 hours. On the day of the experiments, cells were transferred from ND-96 to ND-96’ (ND-96 without gentamicin and sodium pyruvate) containing 8 μM insulin to promote PI3 kinase activity and up-regulate PI(3,4,5)P_3_ levels.

A Leica DM IRBE inverted microscope with a Leica HC Pl APO 20×/0.7 fluorescence objective was used with a Dagan CA-1B amplifier and illuminated with a Lumen Dynamics X-Cite XLED1 light source. Fluorescence was measured with a ThorLabs photomultiplier tube (PMT). The amplifier and light-emitting diode were controlled by a Digidata-1440A board and Axon™ pClamp™ 10.7 software package (Molecular Devices). For the FRET experiments, light was filtered through a HQ436/20 excitation filter and directed to the objective with a 455LP dichroic (Chroma). The microscope cube did not contain an emission filter, because the ThorLabs PMT module contains its own cube. Thus, the emitted light was filtered before the PMTs with a 510-nm dichroic, an HQ480/40 emission filter for CFP, and an HQ535/30 emission filter for YFP (Chroma). For the gPLC experiments, light was filtered through a HQ470/40 excitation ?lter, an HQ525/50 emission ?lter and a Q496LP dichroic (Chroma). The voltage protocol consisted of steps from -100 to 180 mV in irregular increments. The length of the voltage pulse was 1.5-2 s for Xl-VSP1 and varied for Xl-VSP2, 2-10 s for fTAPP, and fPLC, and 3 s for gPLC. Rest periods of 1-3 min between voltage steps were used to allow the cell to recover depleted PIP concentrations before the next voltage step. The resulting FRET or fluorescence was then plotted versus the voltage to generate the fluorescence versus voltage relationship.

### Data analysis

For the FRET experiments, Axon™ Clampfit™ 10.7 (Molecular Devices) was used to calculate the FRET ratio. For the FRET or fluorescence increases (fTAPP and gPLC), the ΔF/F, (*F _after_-F _before_*)/*F _before_*), was calculated from the pre-pulse baseline to the max signal increase. For the fPLC FRET decrease, the ΔF/F was calculated from the pre-pulse baseline to the max FRET decrease. For the fTAPP FRET decrease, the ΔF/F was calculated from the max FRET increase to the max FRET decrease. Calculations were done in Excel (Microsoft) or IGOR Pro (WaveMetrics) and data was plotted in IGOR Pro. When bleaching was observed, it was corrected by fitting to a line and dividing the values by the fit to obtain the final, bleaching-corrected signal. Final voltage dependent curves were plotted with change in FRET or fluorescence on the Y-axis and voltage on the X-axis. The data were fit to single Boltzmann equations. Activity assays were repeated on oocytes extracted on different days from at least three different frogs (biological replicates) until data from a minimum of 8 oocytes were acquired and analyzed. Error bars indicate standard error of the mean (SEM). IHC images were processed with NIS-Elements (Nikon). Fluorescence signals from anti-VSP and matching secondary-alone control images were processed in parallel while applying the same post-image processing parameters in each case. Figures were made either in Illustrator or Photoshop 2018 (Adobe).

## Acknowledgements

We thank Y. Okamura (Osaka University, Osaka, Japan) for kindly providing the Ci-VSP and Dr-VSP cDNAs, E.Y. Isacoff (University of California, Berkeley) for providing the fPLC and fTAPP and T. Meyer for providing the gPLC. We also thank D. Rashid and M. Chaverra for technical assistance and advice.

## Author contributions

Ratzan and S.C. Kohout conceived and designed the experiments. W. Ratzan, S.E. Killian, V. Rayaprolu, R. Bradley and S.C. Kohout collected and analyzed data. W. Ratzan, V. Rayaprolu and S.C. Kohout wrote the manuscript. All authors approved the final version of the manuscript.

## Supporting Information

**Table S1:** Voltage dependence of activity.

| VSP |  | fTAPP |  | fPLC |  | gPLC |  |
| --- | --- | --- | --- | --- | --- | --- | --- |
|  | n | V<sub>1/2</sub> up | V<sub>1/2</sub> down | n | V<sub>1/2</sub> up | n | V<sub>1/2</sub> down |
| XI-VSP1 | 10 | 55 ± 2 | 97 ± 3 | 11 | 98 ± 5 | 11 | 77 ± 3 |
| XI-VSP2 | 11 | 92 ± 8 | n/a | 11 | 83 ± 6 | 9 | 118 ± 3 |

**Fig S1:**
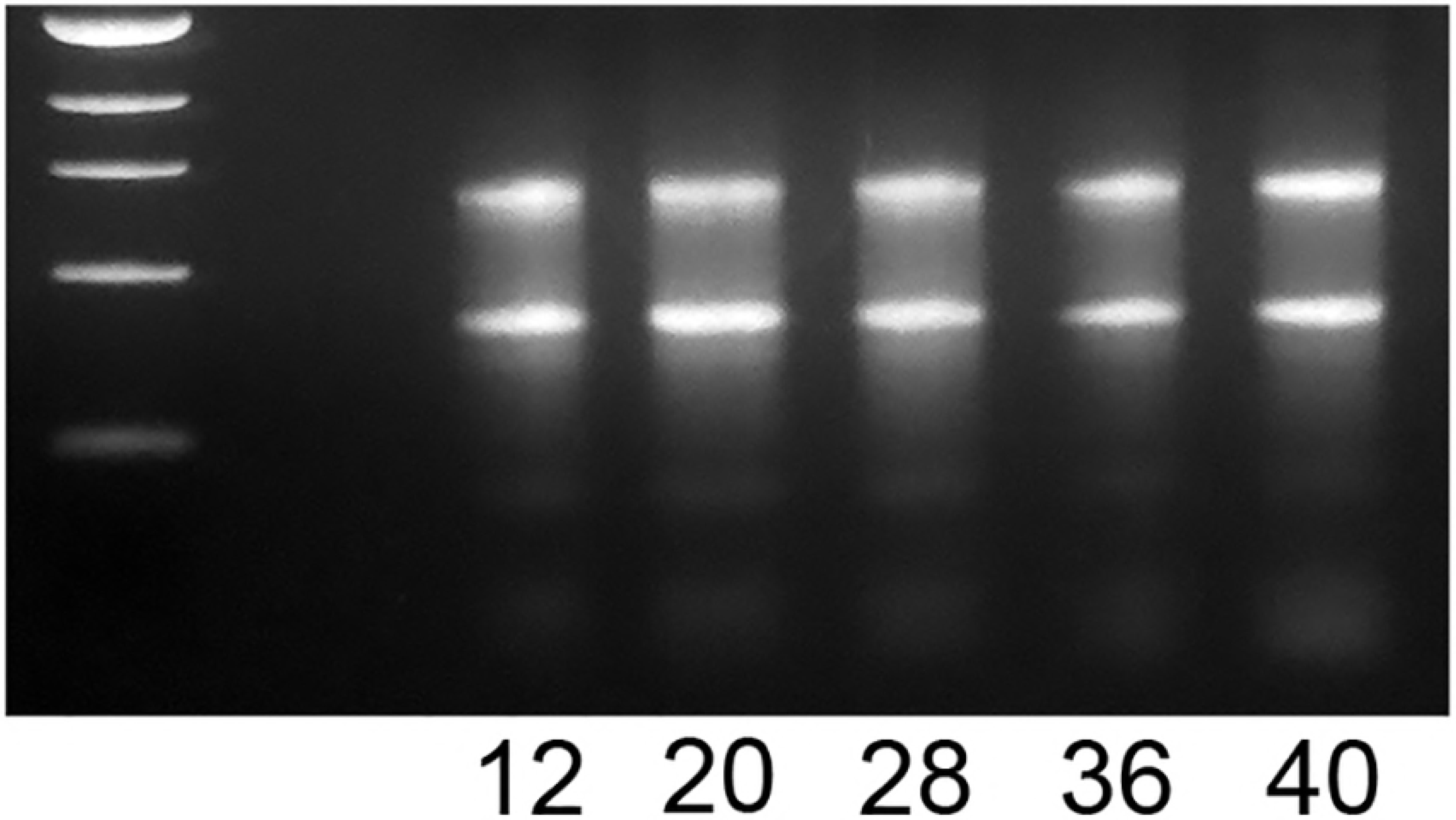
Embryo total RNAs used for sqRT-PCR of Xl-VSPs (NF stages 12-40). RNA concentrations were determined by Nanodrop spectrophotometry and 1 μg of RNA was run on a 2% agarose gel stained with ethidium bromide. The prominent bands seen here are the 28S and 18S ribosomal RNAs, indicating that the RNAs are not degraded and that approximately equal total RNA was used for subsequent sqRT-PCR analysis.

**Fig S2:**
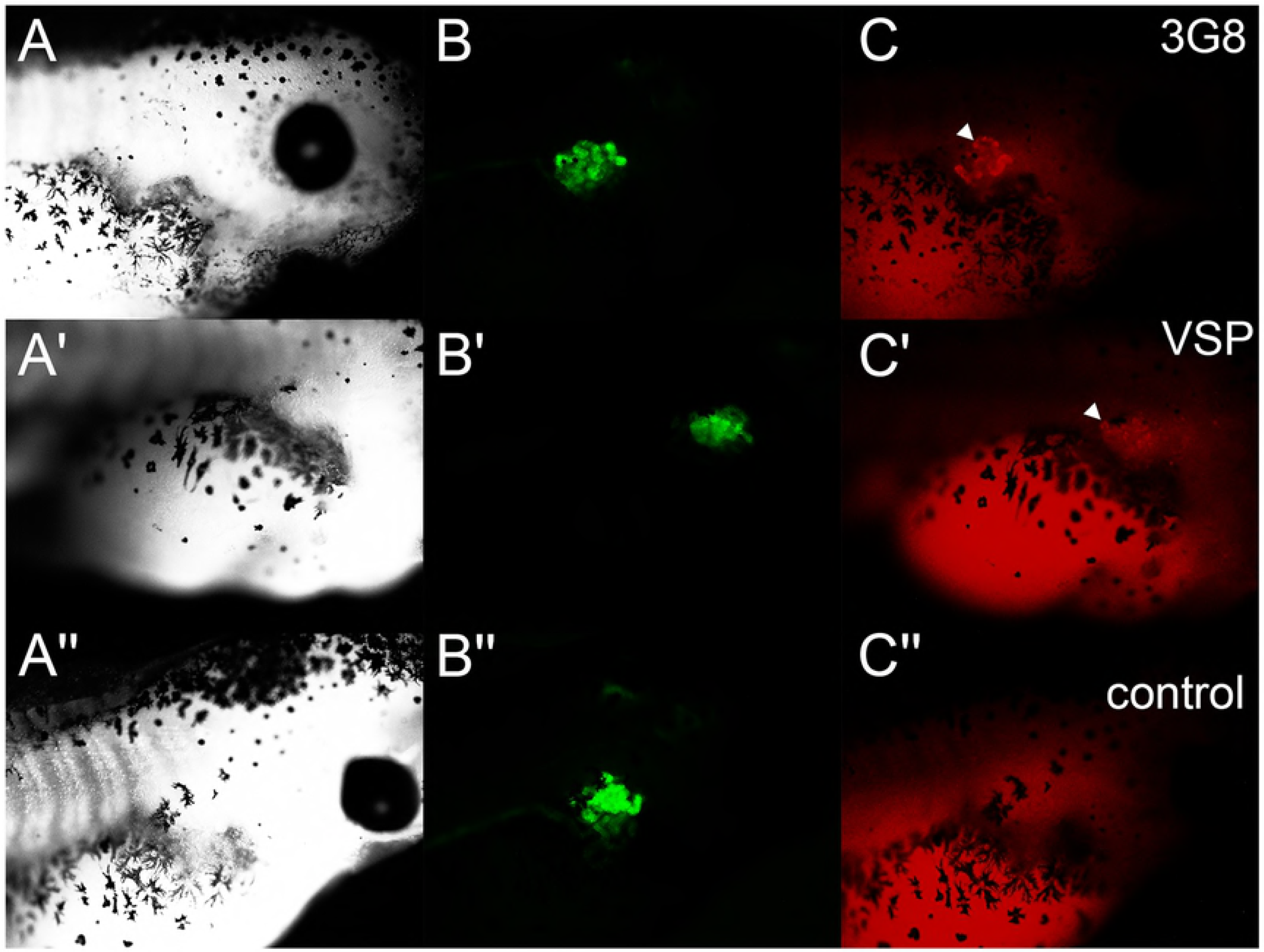
Whole-mount immunohistochemistry (IHC) of NF stage 42 Pax8:GFP embryos. Embryos stained with 3G8 antibody (A-C), anti-VSP (A’-C’) or secondary antibody alone control (A′-C′). The Pax8:GFP embryos show strong GFP expression in the proximal pronephros (B-B′) as confirmed by staining with the proximal pronephros marker 3G8 antibody (C). Xl-VSP staining of tubules is clearly visible (C’) and co-localizes with the GFP, indicating Xl-VSP in the proximal pronephros. Positive staining is marked with arrowheads, whereas no signal above autofluorescence was seen in control animals stained with secondary antibody alone (C′).

